# Characterisation of Gas Vesicles as Cavitation Nuclei for Ultrasound Therapy using Passive Acoustic Mapping

**DOI:** 10.1101/2024.09.24.614126

**Authors:** Cameron A. B. Smith, Avinoam Bar-Zion, Qiang Wu, Dina Malounda, Luca Bau, Eleanor Stride, Mikhail G. Shapiro, Constantin C. Coussios

## Abstract

Genetically encodable gas filled particles known as gas vesicles (GVs) have shown promise as a biomolecular contrast agent for ultrasound imaging and have the potential to be used as cavitation nuclei for ultrasound therapy. In this study, we used passive acoustic mapping techniques to characterize GV-seeded cavitation, utilizing 0.5 and 1.6 MHz ultrasound over peak rarefactional pressures ranging from 100 to 2200 kPa. We found that GVs produce cavitation for the duration of the first applied pulse, up to at least 5000 cycles, but that bubble activity diminishes rapidly over subsequent pulses. At 0.5 MHz the frequency content of cavitation emissions was predominantly broadband in nature, whilst at 1.6 MHz narrowband content at harmonics of the main excitation frequency dominated. Simulations and high-speed camera imaging suggest that the received cavitation emissions come not from individual GVs but instead from the coalescence of GV-released gas into larger bubbles during the applied ultrasound pulse. These results will aid the future development of GVs as cavitation nuclei in ultrasound therapy.

---

Ultrasound-induced cavitation has been shown to have great potential as a therapeutic tool in a wide range of applications. Cavitation has been shown to be effective in temporarily permeabilising the blood-brain barrier^1–3^, in transdermal delivery^4^, and introducing material into individual cells^5–7^. Cavitation has also shown to be a useful means of accelerating heating in thermal ablation therapies^8–10^, whilst simultaneously causing direct mechanical damage to target cells^11,12^. For example, this mechanical damage has been shown to be capable both of killing tumour cells and of stimulating the immune system, leading to improved survival outcomes as well as beneficial abscopal effects^13^.

Exogenous nuclei can be utilized as a method of reducing the ultrasound energy required to generate significant levels of cavitation for use in therapies. A wide variety of particles have been explored as nuclei. For example, coated gas microbubbles have been extensively utilized as contrast agents in ultrasound diagnostic imaging, and have been clinically approved for echocardiography^14^ and microvasculature doppler imaging^15^, and have more recently shown promise as cavitation agents for use in blood brain barrier opening^1–3^and cancer therapy^16^. Solid gas-stabilizing particles^17,18^ have shown promise in enabling persistent cavitation over tens of minutes both within vascular and extravascular spaces ^11,12^.

Gas Vesicles (GVs) are protein-shelled gas filled particles^19^ that can be expressed by genetically engineered bacteria^20^ and mammalian^21–23^ cells. GVs have previously been characterized for high frequency short pulses for the purpose of ultrasound imaging^20,24–26^. Amplitude modulation imaging^27^ capitalises on the buckling properties of GVs to generate non-linear signals which can be exploited to achieve improved signal to background ratio^28,29^. BURST imaging^30^ takes advantage of the collapsibility of GVs with higher pressure pulses by unmixing the unique transient signal associated with GV collapse from the linear scattering background, resulting in single-cell sensitivity.

GV-producing cells are capable of infiltrating the cores of tumours^22,31^, and are capable of being engineered such that they only express the GVs under certain conditions^30,32^. This allows for a high degree of molecular specificity, as well as allowing the GVs to populate regions of tumours which are traditionally the hardest to reach with conventional cavitation nuclei^22,31^. GVs therefore occupy an interesting potential niche as a cavitation nucleation tool for ultrasound therapy. An initial study exploring their use as nucleation points for mechanical ablation has shown promise^31^. In this work we conduct a thorough characterization of the cavitation dynamics of GVs to guide future biomolecular and ultrasound parameters for GV-enabled cavitation-based therapy.

To characterize the GVs, we used Passive Acoustic Mapping (PAM), a source coherence passive beamforming process^33^ which can be implemented in the time or frequency domain^34^. PAM captures the cavitation emissions emanating from a specific location with high SNR. We used recently developed robust Capon beamforming (RCB) PAM, which utilizes optimized beamformer weightings to reduce artefacts, further improving the SNR^35^.

The experimental set-up (figure 1) we used to characterise the GVs comprised a single spherically focussed therapeutic transducer of 64 mm active diameter and focal length 62.6 mm (H-107 D SN12; Sonic Concepts, Bothell, WA, USA) generating ultrasound at a center frequency of either 0.5 or 1.6 MHz and peak rarefactional pressures ranging from 0-2.2 MPa. Samples were suspended in 0.5 ml volumes in 2 ml microcentrifuge tubes (Eppendorf LoBind, 2 mL; Sigma Aldrich, Hamburg, Germany). The therapeutic transducer had a central rectangular cutout of dimensions 45-18mm to enable positioning of a co-axial linear array for passive acoustic mapping of cavitation. Cavitation signals generated were received using two co-planar L7-4 linear arrays (L7-4, Verasonics, Kirkland, WA, USA), one positioned inside and coaxially with the therapeutic transducer, the second at 90 degrees to the therapeutic transducer acoustic axis, which were calibrated using a wire scatter^36^. These signals were recorded using a programmable ultrasound engine (Vantage 256, Verasonics) and then beamformed using the RCB-PAM algorithm in a 10 mm lateral by 15 mm axial region of interest, with a pixel size of 0.4 mm square, ε = 5. GV’s were isolated form *Anabaena flos-aquae* as described previously^37^.

**FIG. 1.**
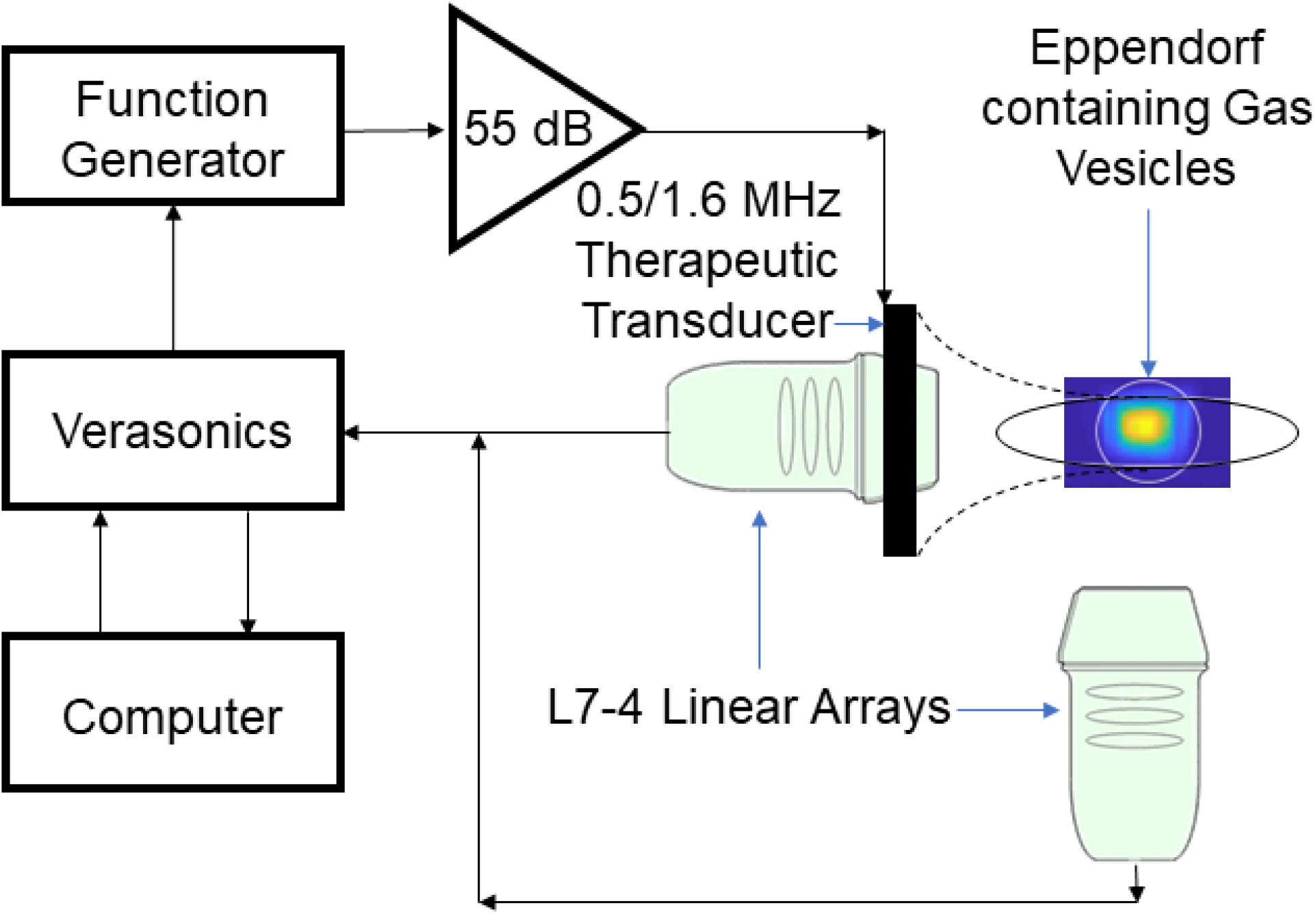
Experimental set-up, consisting of a therapeutic transducer transmitting either its central frequency 0.5 MHz or at the third harmonic 1.6 MHz at a range of ultrasound parameters, focused on a microcentrifuge tube containing anabaena gas vesicles of a range of concentrations. The cavitation emissions are then received using two perpendicular co-planar L7-4 linear arrays for later post-processing.

First, we explored the persistence of the generated cavitation. We discovered that cavitation emissions can be detected for the duration of the applied pulse up to 5000 cycles (figures 2a,b) with a pulse repetition frequency (PRF) of 0.25 Hz. However, upon subsequent pulses the cavitation levels are greatly diminished, with very little cavitation generated after 4 pulses (figures 2c,d). We also found this to be true at lower pressures, down to 200 kPa peak negative pressure. This suggests that if the intention is to maximize cavitation activity, one should focus on long pulses with a PRF set to allow for recirculation of GVs in the acoustic focus, assuming that recirculation is feasible for the required application. Due to this low persistence for all further cavitation analysis in this characterization study we utilised only the signals emitted during the first cavitation pulse.

**FIG. 2.**
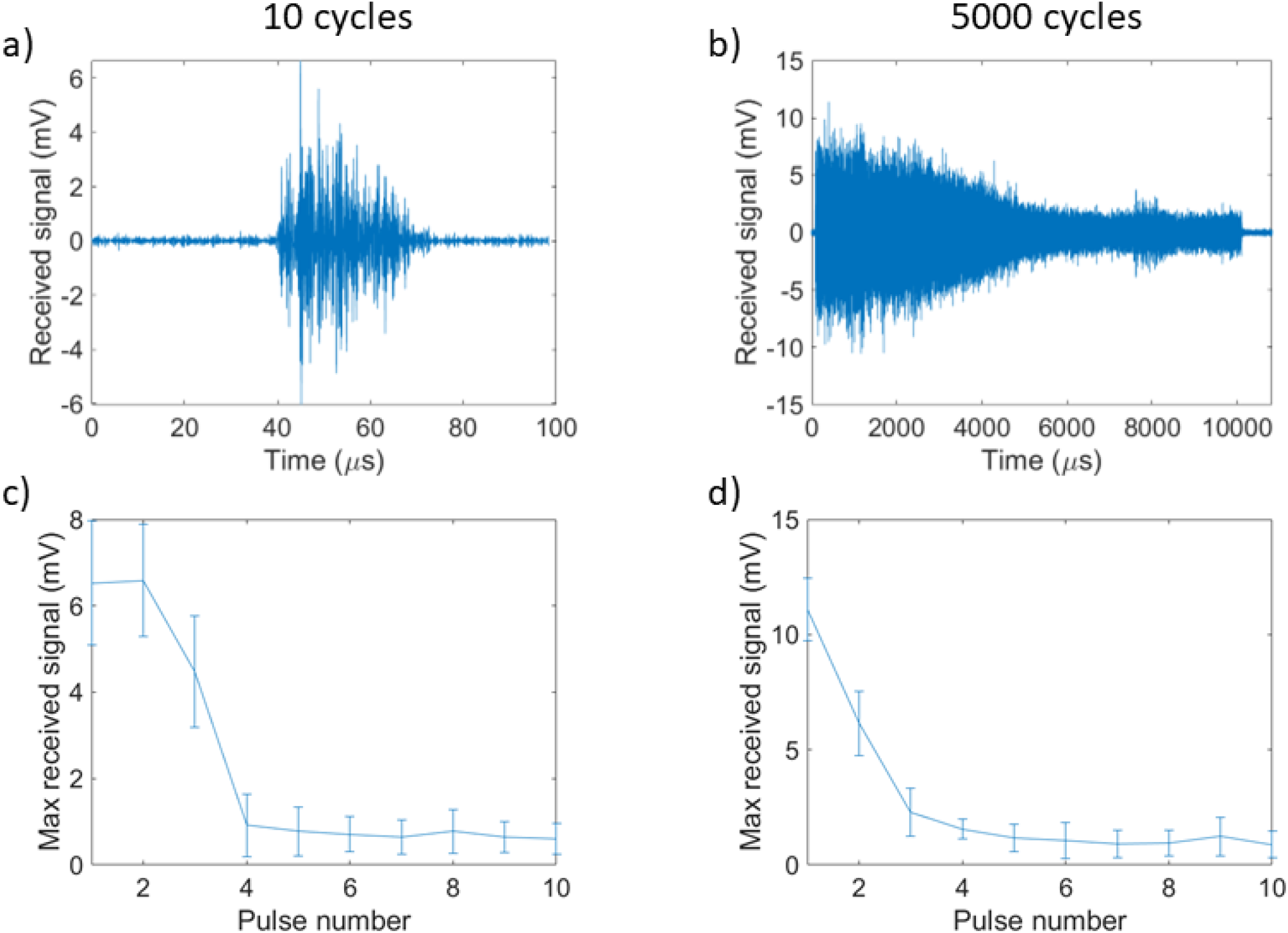
Representative example of the persistence of cavitiation over the course of the first transmitted pulse for 10 (a) and 5000 (b) cycles, 0.5 MHz, 1 MPa, 8×10^10^ particles/mL, 0.25 Hz PRF. Maximum received signal over subsequent therapeutic ultrasound pulses, averaged over 10 samples with error bars representing the standard deviation for 10 (c) and 5000 (d) cycles.

Next, we examined the influence of applied pressure, both on GVs and buffered saline (PBS) control samples.

Cavitation emissions from the GVs were significantly higher than from PBS, with emission levels increasing with increasing pressure as expected for both 0.5 and 1.6 MHz (figure 3), consistent with previous results^31^. We separated the cavitation emissions into their broadband and harmonic components using a perfect comb filter in the frequency domain, with the filter multiplying the power spectral density of emissions by 0 at the harmonic and ultra-harmonic frequencies ± 25 kHz, and by 1 everywhere else, resulting in the removal of harmonics and any broadband emissions that happened to overlap with the 50kHz bands of the comb filter.

**FIG. 3.**
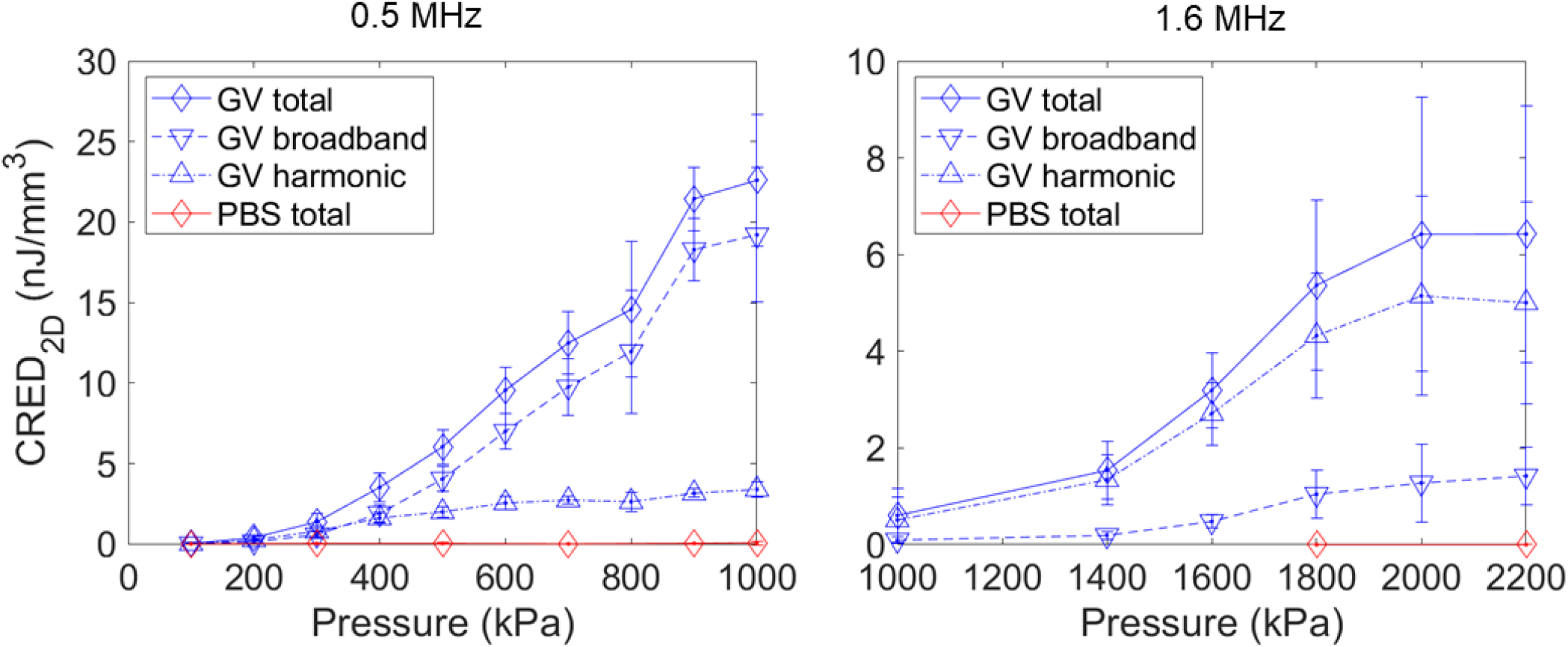
Average Cavitation Radiated Energy Density (CRED) (n=10) split into total, broadband, and harmonic emissions over a range of applied peak negative pressures at 0.5 and 1.6 MHz incident frequency, 1.4×10^10^ particles/mL, with error bars representing standard deviation.

The broadband energy lost by the filtered-out broadband signal was recovered by scaling the result by the number of 1s and 0s in the filter divided by the number of 1s in the filter, with the length of the filter being determined by the usable bandwidth of the transducer. With the full energy of the broadband emissions recovered, the total energy of harmonic emissions were then estimated by subtracting the broadband energy from the total emissions energy. As can be seen in figure 3, both the broadband and harmonic energies grew with increasing applied pressure, with the majority of the emissions being broadband in the case of 0.5 MHz, and harmonic in the case of 1.6 MHz, even at comparable cavitation indexes^38^. The total emissions are four times stronger in the 0.5 MHz case than the 1.6 MHz case, with this being driven almost entirely by the difference in broadband emissions.

To gain a better understanding of the GV behavior during the applied therapeutic pulse, we generated spectrograms to see the distribution of emissions over the duration of the pulse, which we repeated at a range of particle concentrations (figure 4). We generated these spectrograms by averaging the spectrograms of the beamformed signals at the center of the tube over 10 samples to improve the SNR. In the case of the 500 kHz-stimulated emissions some interesting temporal dynamics could be observed. Initially a quick build up can be observed as the harmonics and then the broadband emissions rise, until after a certain point the broadband starts to reduce and only the harmonics remain, which slowly decrease in intensity until the end of the applied acoustic pulse. The broadband signal appears to continue for a longer period of time at higher GV concentrations. In the case of the 1.6 MHz-stimulated emissions consistent harmonic and ultra-harmonic emissions can be seen at all tested concentrations.

**FIG. 4.**
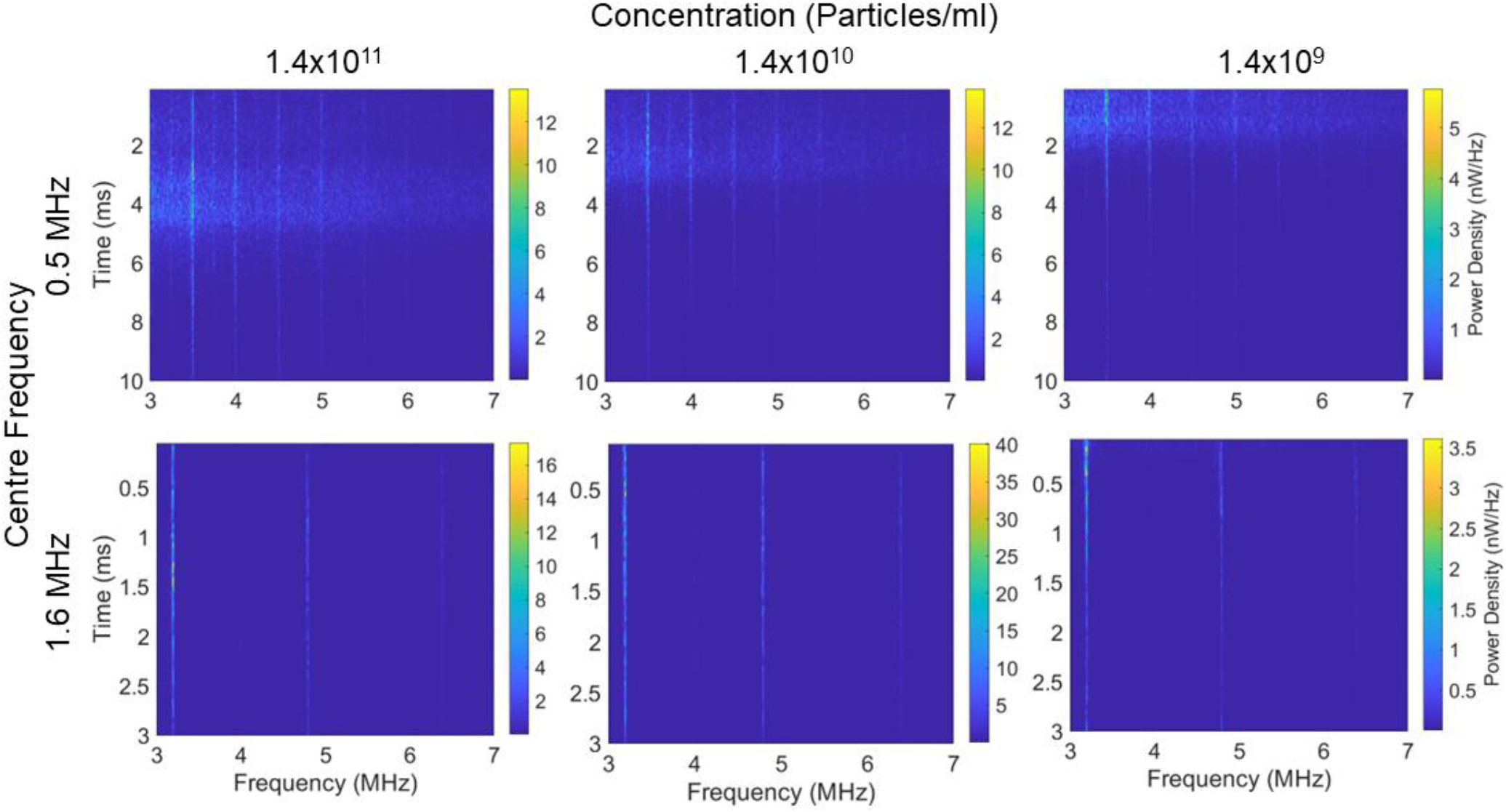
Average spectrograms over 5 samples showing the frequency distribution of cavitation emissions over the pulse duration for a range of particle concentrations and incident frequencies when exposed to a 5000 cycle pulse of 1 MPa in the case of 0.5 MHz and 2 MPa in the case of 1.6 MHz.

To better understand these emissions, we conducted a series of bubble simulations. Previously it has been theorized that under high pressure pulses the GV is quickly destroyed, releasing a free bubble of the gas trapped inside^30,31^, with the subsequent radiated acoustic emissions being those which are generated from the free bubble. We used a Keller-Miksis model, as it has been previously shown to accurately predict both broadband and harmonic acoustic emissions that are in both qualitative and quantitative agreement with experimental measurements^39^. Our parameters can be seen in table 1. We used a range of R0 values to explore bubbles with different rest sizes (figure 5b). Of particular interest is the initial stages of insonation, figure 5a shows the experimental spectrogram from figure 4b cropped to focus on the first 100 *μ*s, with t = 0 defined as the timepoint accounting for the time of flight from the therapeutic transducer to the center of the microcentrifuge tube and back to the linear array, corresponding to the time at which one would expect to receive the start of the cavitation signal. It can be seen here that there appears to be a small delay before we begin to see the cavitation signal, the harmonic components come in, and the broadband component also grows over time.

**TABLE I.**
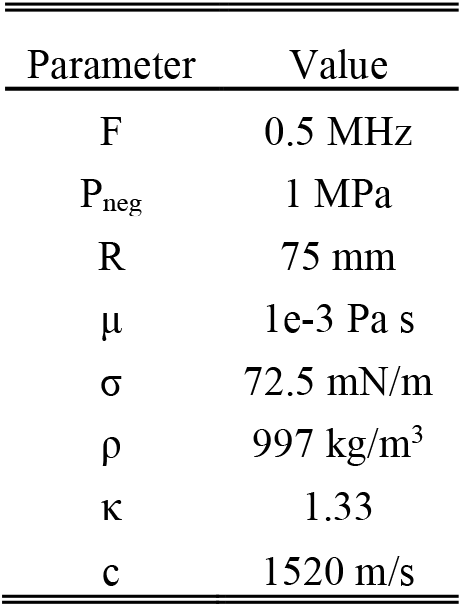
Keller-Miksis Parameters.

| Parameter | Value |
| --- | --- |
| F | 0.5 MHz |
| $P_{\text{neg}}$ | 1 MPa |
| R | 75 mm |
| $\mu$ | 1e-3 Pa s |
| $\sigma$ | 72.5 mN/m |
| $\rho$ | 997 kg/m<sup>3</sup> |
| $\kappa$ | 1.33 |
| c | 1520 m/s |

**FIG. 5.**
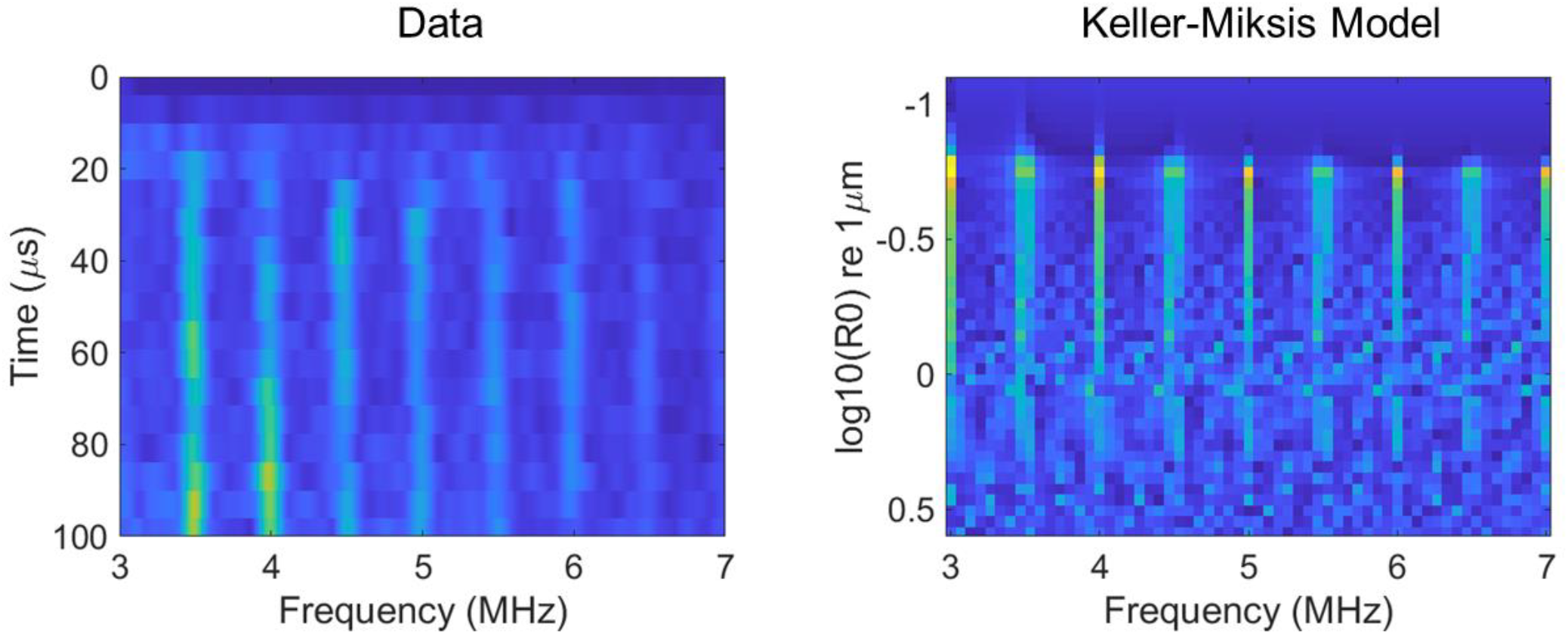
Average spectrogram (n=5) showing the frequency distribution of cavitation emissions over the pulse duration for 0.5 MHz and 1.4×10^10^ particle concentration, cropped to more clearly show the cavitation emissions during the first 50 transmitted cycles, Time = 0 corresponds to the timepoint at which cavitation emissions are expected to be received from the center of the Eppendorf (left). On the right the results for a Keller-Miksis bubble simulation showing the frequency distribution of cavitation emissions from single bubbles at a range of bubble resting sizes (R0) when exposed to the same incident ultrasound parameters and receiving array bandwidth.

A single *Anabaena* GV has an approximate size of a 85 nm diameter cylinder with a length in the range of 200-800 nm (average 519 nm)^37,40^. A sphere of equivalent volume would have a radius of roughly 90 nm. In practice the radius of the equivalent bubble would likely be smaller than this given that the surface tension experienced by the gas will be greater without the presence of the protein shell, but we shall take 90 nm as a conservative estimate. It can be seen by the Keller-Miksis simulation that a bubble of this size does not give significant cavitation emissions when exposed to the ultrasound parameters used in this study; however, a larger bubble would. We propose, therefore, that the bubble emissions we record are not due to single GVs but instead originate from larger bubbles generated by coalescence of the bubbles released from GVs. Our hypothesis is that the initial ultrasound cycles collapse the protein shells, releasing small gas bubbles without measurable cavitation emissions, and over the first 15 *μ*s this gas coalesces into larger bubbles which are capable of producing harmonic and broadband emissions under continued acoustic excitation. At later timepoints seen in figure 4 the less stable inertially cavitating bubbles eventually dissipate, forming larger, stable bubbles which emit predominantly harmonic emissions that gradually become smaller as gas diffuses slowly into the surrounding medium. The time it takes for this to occur is longer the higher the total gas volume, explaining why the period over which broadband emissions are emitted is longer at the higher GV concentrations in figures 4a-c.

To explore this theory further we modified the set-up (figure 6) to allow for the addition of a high-speed camera (HPV-X2, Shimadzu, Japan) running at 1 million frames per second, thereby allowing for simultaneous optical and acoustic imaging. A range of ultrasound parameters and particle concentrations were used. 50 cycle pulses were used so that the entirety of the cavitation time could be recorded optically and acoustically. It is worth noting that this new set-up has a significantly lower volume of sample within the focal region, with a polyethylene tube of 180 *μ*m inner diameter and 10 *μ*m wall thickness (Advanced Polymers, Salem NH, USA) being targeted instead of a 0.5 ml volume in a microcentrifuge tube. Within this new dataset a tight threshold for cavitation emissions can be seen as demonstrated in figure 7, where the percentage of repeat samples (n=30) which had at any point in time an SNR > 10 is shown for a range of free-field pressures and concentrations. While using a concentration of 1.2×10^10^ particles per ml at a pressure of 1 MPa 90% of the samples gave cavitation emissions, if the pressure was reduced to 0.8 MPa or if the particle concentration was reduced by a factor of 1.5 then none of the 30 samples produced cavitation emissions upon exposure. Note that at this concentration the average distance between particles is approximately 5 *μ*m, with hundreds of thousands of GVs within the acoustic field of view. If we look at our optical field of view (figure 8a), with t = 0 being the frame immediately before the ultrasound arrives in the field of view, we can see that the background within the tube becomes brighter over the first 4 μs, then a few bubbles which are large enough to be viewed optically appear, coalesce and oscillate for the duration of the 100 μs long pulse, and then dissipate upon the conclusion of the pulse, with the background remaining at the brighter level. Figure 8b shows a plot of the percent change in background intensity across a region away from oscillating bubbles. We theorize that this background intensity corresponds to light scattering by the GVs, that what we observe optically is the quick destruction of all the GVs in the focal region, releasing the gas within, this gas then either quickly dissolves into the surrounding media or, if there happens to be within the region enough gas to quickly coalesce, it does so. The gas quickly coalesces into a micron-sized bubble to oscillate and release the recorded acoustic emissions. This would explain the sharp threshold we see, the delay in received emissions, and how the change in optical intensity between the start and the end scales with concentration.

**FIG. 6.**
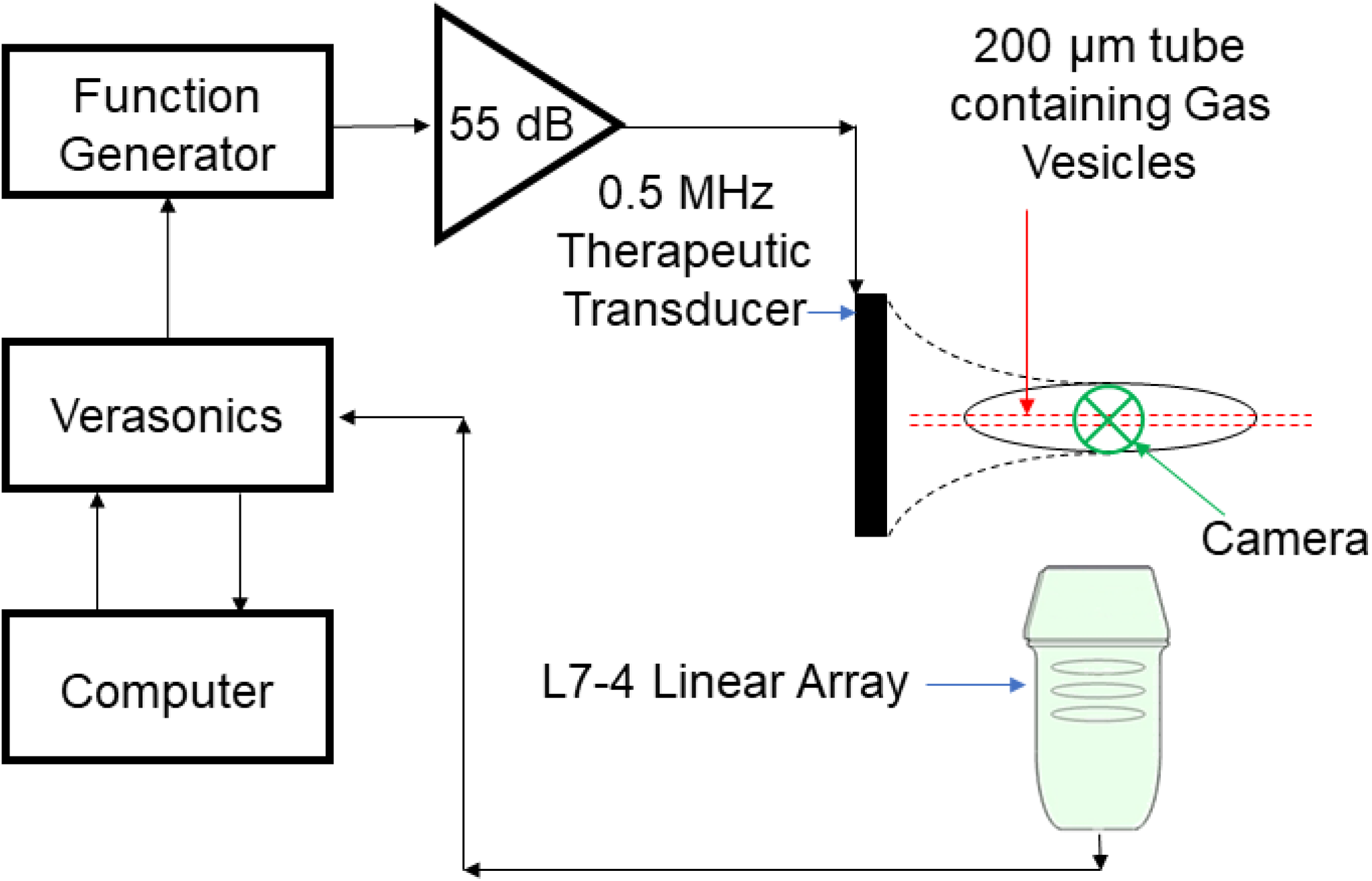
Experimental set-up, consisting of a therapeutic transducer transmitting its central frequency 0.5 MHz at a range of ultrasound parameters, focused on a 200 μm diameter polyethylene tube containing gas vesicles at a range of concentrations. The cavitation emissions are then received using an L7-4 linear array for later post-processing.

**FIG. 7.**
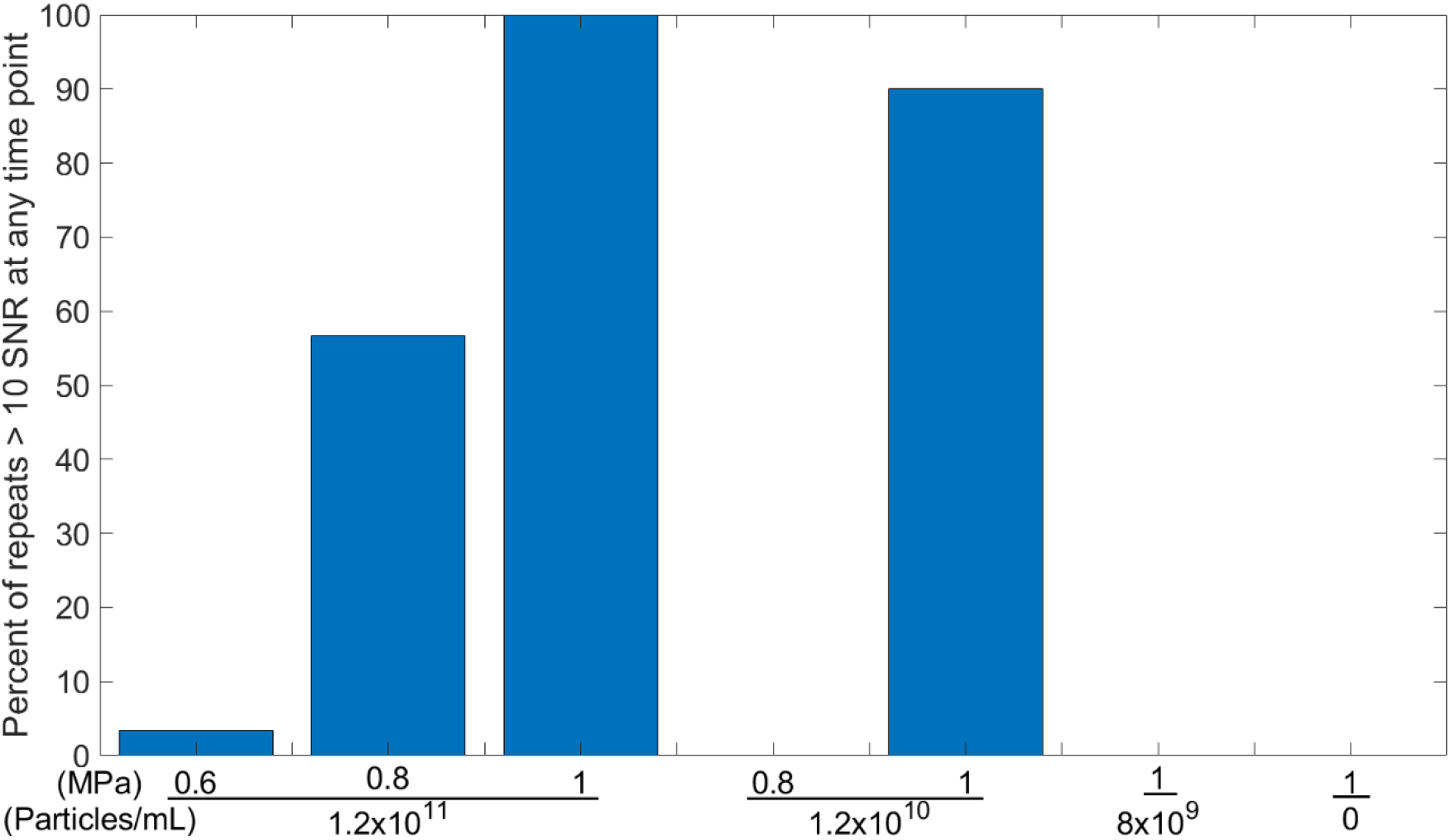
Percentage of samples (n=30) that had at any timepoint a notable cavitation signal (defined as SNR > 10) at a range of peak negative pressures and particle concentrations.

**FIG. 8.**
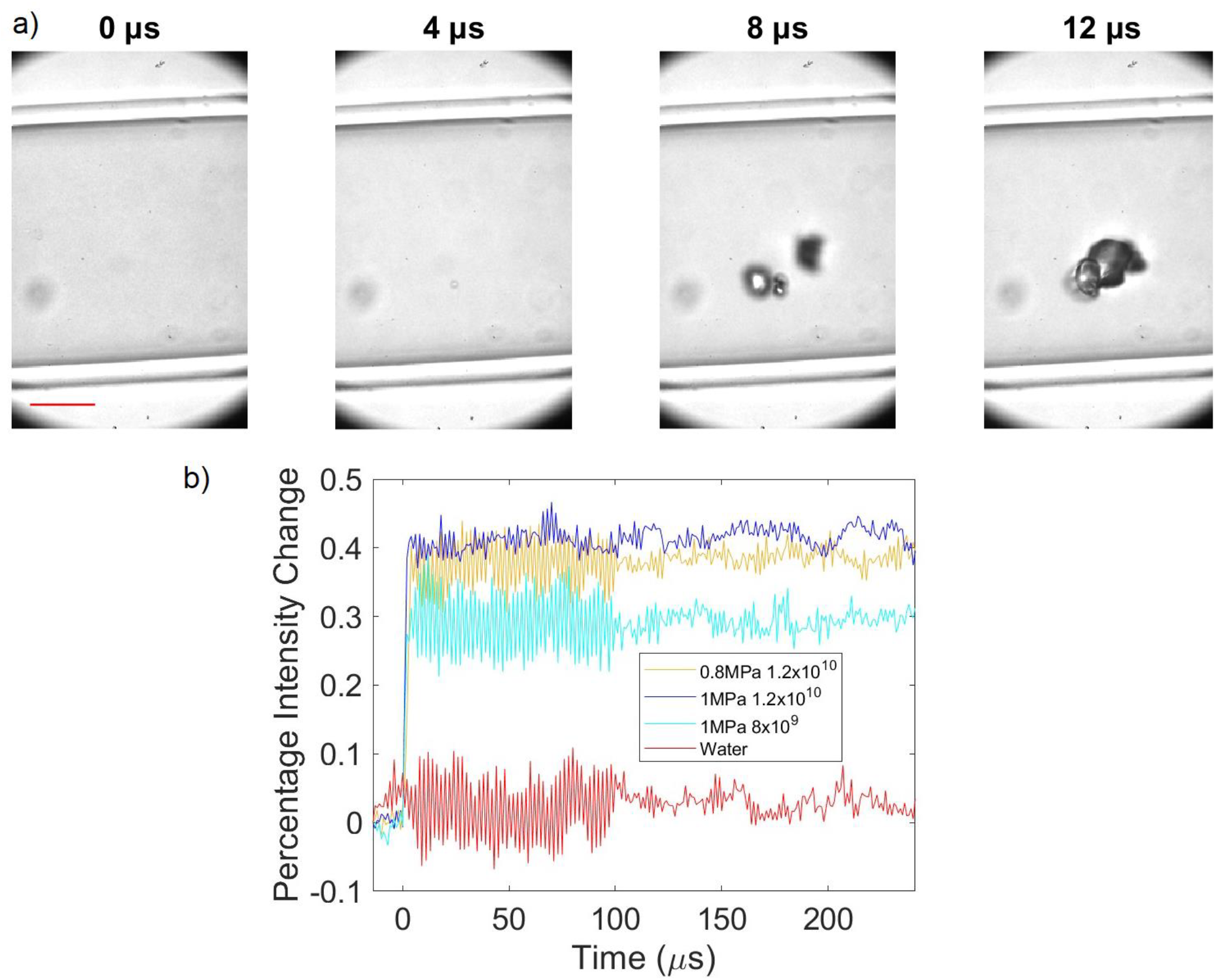
**(a)** Frames from a representative high-speed video at 0.8 MPa and 1.2×10^11^ particles/ml with Time = 0 corresponds to the time at which the incident therapeutic pulse reaches the center of the tube, scale bar = 50 *μ*m. (b) Percentage intensity change over time in the optical image in a region of interest outside of the bubbles but within the tube at a range of applied pressures and particle concentrations.

In conclusion, GVs are capable of acting as cavitation nuclei while using therapeutic pulses, though their irreversible collapse and gas release requires the use of a low number of longer pulses in order to maximize the energy output. Throughout the range of pressures investigated, GVs underwent predominantly inertial cavitation when exposed to 0.5 MHz ultrasound and non-inertial cavitation at 1.6 MHz. From the combined simulations and experimental evidence, it appears that the cavitation emissions detected come not from individual GVs but instead from micron-scaled bubbles formed by the coalescence of gas released from destroyed GVs during the emission pulse. A sharp cavitation threshold was discovered at low sample volumes both in terms of applied acoustic pressure and GV concentration, which gives further credence to the hypothesis that nanobubble release and coalescence is a necessary pre-condition to the formation of acoustically excitable cavities at therapeutic ultrasound frequencies and pressures. These results will aid in the future use of GVs as cavitation nuclei for ultrasound therapy by informing the acoustic parameters driving desired cavitation behaviour, the interpretation of mechanical effects caused by GV-seeded cavitation, and the engineering of GVs and GV-expressing cells achieve desired cavitation profiles.

## Acknowledgements

The authors gratefully acknowledge funding from the International Human Frontier Science Program Organization (grant LT0036/2022-L) and the National Institutes of Health (R01-EB018975 to M.G.S.). Related research in the Shapiro laboratory is supported by the David and Lucille Packard Foundation and the Chan Zuckerberg Initiative. M.G.S. Is an Investigator of the Howard Hughes Medical Institute.

## REFERENCES

(1) Arvanitis, C. D.; Ferraro, G. B.; Jain, R. K. The Blood–Brain Barrier and Blood–Tumour Barrier in Brain Tumours and Metastases. Nature Reviews Cancer 2020, 20 (1), 26–41. 10.1038/s41568-019-0205-x.

(2) Hynynen, K.; McDannold, N.; Vykhodtseva, N.; Jolesz, F. A. Noninvasive MR Imaging–Guided Focal Opening of the Blood-Brain Barrier in Rabbits. Radiology 2001, 220 (3), 640–646. 10.1148/radiol.2202001804.

(3) Mcmahon, D.; Reilly, M. A. O.; Hynynen, K. Therapeutic Agent Delivery Across the Blood–Brain Barrier Using Focused Ultrasound. Annual Review of Biomedical Engineering 2021, 89–113.

(4) Bhatnagar, S.; Kwan, J. J.; Shah, A. R.; Coussios, C. C.; Carlisle, R. C. Exploitation of Sub-Micron Cavitation Nuclei to Enhance Ultrasound-Mediated Transdermal Transport and Penetration of Vaccines. Journal of Controlled Release 2016, 238, 22–30. 10.1016/j.jconrel.2016.07.016.

(5) Lentacker, I.; De Cock, I.; Deckers, R.; De Smedt, S. C.; Moonen, C. T. W. Understanding Ultrasound Induced Sonoporation: Definitions and Underlying Mechanisms. Advanced Drug Delivery Reviews 2014, 72, 49–64. 10.1016/j.addr.2013.11.008.

(6) Okada, K.; Kudo, N.; Niwa, K.; Yamamoto, K. A Basic Study on Sonoporation with Microbubbles Exposed to Pulsed Ultrasound. Journal of Medical Ultrasonics 2005, 32 (1), 3–11. 10.1007/s10396-005-0031-5.

(7) Beekers, I.; Vegter, M.; Lattwein, K. R.; Mastik, F.; Beurskens, R.; van der Steen, A. F. W.; de Jong, N.; Verweij, M. D.; Kooiman, K. Opening of Endothelial Cell–Cell Contacts Due to Sonoporation. Journal of Controlled Release 2020, 322 (March), 426–438. 10.1016/j.jconrel.2020.03.038.

(8) Den Brok, M. H. M. G. M.; Sutmuller, R. P. M.; Van Der Voort, R.; Bennink, E. J.; Figdor, C. G.; Ruers, T. J. M.; Adema, G. J. In Situ Tumor Ablation Creates an Antigen Source for the Generation of Antitumor Immunity. Cancer Research 2004, 64 (11), 4024– 4029. 10.1158/0008-5472.CAN-03-3949.

(9) Zhang, L.; Wang, Z.-B. High-Intensity Focused Ultrasound Tumor Ablation: Review of Ten Years of Clinical Experience. Frontiers of Medicine in China 2010, 4 (3), 294–302. 10.1007/s11684-010-0092-8.

(10) Huang, H.; Ran, J.; Xiao, Z.; Ou, L.; Li, X.; Xu, J.; Wang, Q.; Wang, Z.; Li, F. Reasons for Different Therapeutic Effects of High-Intensity Focused Ultrasound Ablation on Excised Uterine Fibroids with Different Signal Intensities on T2-Weighted MRI: A Study of Histopathological Characteristics. International Journal of Hyperthermia 2019, 0 (0), 1– 8. 10.1080/02656736.2019.1592242.

(11) Smith, C. A. B.; Coussios, C. C. Spatiotemporal Assessment of the Cellular Safety of Cavitation-Based Therapies by Passive Acoustic Mapping. Ultrasound in Medicine and Biology 2020, 46 (5), 1235–1243. 10.1016/j.ultrasmedbio.2020.01.009.

(12) Smith, C. A. B.; Coussios, C. C. Spatiotemporal Assessment of the Cellular Safety of Cavitation-Based Therapies by Passive Acoustic Mapping. He 24th European symposium on Ultrasound Contrast Imaging 194–195.

(13) Qu, S.; Worlikar, T.; Felsted, A. E.; Ganguly, A.;Beems, M. V.; Hubbard, R.; Pepple, A. L.; Kevelin, A. A.; Garavaglia, H.; Dib, J.; Toma, M.; Huang, H.;Tsung, A.; Xu, Z.; Cho, C. S. Non-ThermalHistotripsy Tumor Ablation Promotes AbscopalImmune Responses That Enhance Cancer Immunotherapy. Journal for ImmunoTherapy of Cancer 2020, 8 (1), 1–12. 10.1136/jitc-2019-000200.

(14) Galema, T. W.; Geleijnse, M. L.; Vletter, W. B.; deLaat, L.; Michels, M.; ten Cate, F. J. Clinical Usefulness of SonoVue Contrast Echocardiography: The Thoraxcentre Experience. Neth Heart J 2007, 15 (2), 55–60.

(15) Mitchell, W. K.; Phillips, B. E.; Williams, J. P.;Rankin, D.; Smith, K.; Lund, J. N.; Atherton, P. J. Development of a New Sonovue™ Contrast-Enhanced Ultrasound Approach Reveals Temporal and Age-Related Features of Muscle Microvascular Responses to Feeding. Physiological Reports 2013, 1 (5). 10.1002/phy2.119.

(16) Carlisle, R.; Choi, J.; Bazan-Peregrino, M.; Laga, R.;Subr, V.; Kostka, L.; Ulbrich, K.; Coussios, C. C.;Seymour, L. W. Enhanced Tumor Uptake and Penetration of Virotherapy Using Polymer Stealthing and Focused Ultrasound. Journal of the National Cancer Institute 2013, 105 (22), 1701–1710.10.1093/jnci/djt305.

(17) Kwan, J. J.; Myers, R.; Coviello, C. M.; Graham, S.M.; Shah, A. R.; Stride, E.; Carlisle, R. C.; Coussios, C. C. Ultrasound-Propelled Nanocups for Drug Delivery. Small 2015, 11 (39), 5305–5314.

(18) Montoya Mira, J.; Wu, L.; Sabuncu, S.; Sapre, A.;Civitci, F.; Ibsen, S.; Esener, S.; Yildirim, A. Gas-Stabilizing Sub-100 Nm Mesoporous Silica Nanoparticles for Ultrasound Theranostics. ACS Omega 2020, 5 (38), 24762–24772.10.1021/acsomega.0c03377.

(19) Lu, G. J.; Farhadi, A.; Mukherjee, A.; Shapiro, M. G. Proteins, Air and Water: Reporter Genes for Ultrasound and Magnetic Resonance Imaging. Current Opinion in Chemical Biology 2018, 45, 57–63. 10.1016/j.cbpa.2018.02.011.

(20) Shapiro, M. G.; Goodwill, P. W.; Neogy, A.; Yin, M.;Foster, F. S.; Schaffer, D. V.; Conolly, S. M. BiogenicGas Nanostructures as Ultrasonic MolecularReporters. Nature Nanotechnology 2014, 9 (4), 311–316. 10.1038/nnano.2014.32.

(21) Bourdeau, R. W.; Lee-Gosselin, A.; Lakshmanan, A.;Farhadi, A.; Kumar, S. R.; Nety, S. P.; Shapiro, M. G. Acoustic Reporter Genes for Noninvasive Imaging of Microorganisms in Mammalian Hosts. Nature 2018, 553 (7686), 86–90.10.1038/nature25021.

(22) Hurt, R. C.; Buss, M. T.; Duan, M.; Wong, K.; You, M. Y.; Sawyer, D. P.; Swift, M. B.; Dutka, P.; Barturen-Larrea, P.; Mittelstein, D. R.; Jin, Z.; Abedi, M. H.; Farhadi, A.; Deshpande, R.; Shapiro, M. G. Genomically Mined Acoustic Reporter Genes for Real-Time in Vivo Monitoring of Tumors and Tumor-Homing Bacteria. Nat Biotechnol 2023, 1–13. 10.1038/s41587-022-01581-y.

(23) Farhadi, A.; Ho, G. H.; Sawyer, D. P.; Bourdeau, R. W.; Shapiro, M. G. Ultrasound Imaging of Gene Expression in Mammalian Cells. Science 2019, 365 (6460), 1469–1475. 10.1126/science.aax4804.

(24) Cherin, E.; Melis, J. M.; Bourdeau, R. W.; Yin, M.; Kochmann, D. M.; Foster, F. S.; Shapiro, M. G. Acoustic Behavior of Halobacterium Salinarum Gas Vesicles in the High-Frequency Range: Experiments and Modeling. Ultrasound in Medicine and Biology 2017, 43 (5), 1016–1030. 10.1016/j.ultrasmedbio.2016.12.020.

(25) Yang, Y.; Qiu, Z.; Liu, C.; Huang, Y.; Sun, L.; Dai, J. Acoustic Characterization of Nano Gas Vesicles. 2015 IEEE International Ultrasonics Symposium, IUS 2015 2015, 1–4. 10.1109/ULTSYM.2015.0386.

(26) Zhang, S.; Huang, A.; Bar-Zion, A.; Wang, J.; Mena, O. V.; Shapiro, M. G.; Friend, J. The Vibration Behavior of Sub-Micrometer Gas Vesicles in Response to Acoustic Excitation Determined via Laser Doppler Vibrometry. Advanced Functional Materials 2020, 30 (13), 2000239. 10.1002/adfm.202000239.

(27) Maresca, D.; Sawyer, D. P.; Renaud, G.; Lee-Gosselin, A.; Shapiro, M. G. Nonlinear X-Wave Ultrasound Imaging of Acoustic Biomolecules. Physical Review X 2018, 8 (4), 41002. 10.1103/PhysRevX.8.041002.

(28) Maresca, D.; Lakshmanan, A.; Lee-Gosselin, A.; Melis, J. M.; Ni, Y. L.; Bourdeau, R. W.; Kochmann, D. M.; Shapiro, M. G. Nonlinear Ultrasound Imaging of Nanoscale Acoustic Biomolecules. Applied Physics Letters 2017, 110 (7). 10.1063/1.4976105.

(29) Rabut, C.; Wu, D.; Ling, B.; Jin, Z.; Malounda, D.; Shapiro, M. G. Ultrafast Amplitude Modulation for Molecular and Hemodynamic Ultrasound Imaging. Appl. Phys. Lett. 2021, 118 (24), 244102. 10.1063/5.0050807.

(30) Sawyer, D. P.; Bar-Zion, A.; Farhadi, A.; Shivaei, S.; Ling, B.; Lee-Gosselin, A.; Shapiro, M. G. Ultrasensitive Ultrasound Imaging of Gene Expression with Signal Unmixing. Nature Methods 2021, 18 (8), 945–952. 10.1038/s41592-021-01229-w.

(31) Bar-Zion, A.; Nourmahnad, A.; Mittelstein, D. R.; Shivaei, S.; Yoo, S.; Buss, M. T.; Hurt, R. C.; Malounda, D.; Abedi, M. H.; Lee-Gosselin, A.; Swift, M. B.; Maresca, D.; Shapiro, M. G. Acoustically Triggered Mechanotherapy Using Genetically Encoded Gas Vesicles. Nature Nanotechnology 2021, 33. 10.1038/s41565-021-00971-8.

(32) Lakshmanan, A.; Jin, Z.; Nety, S. P.; Sawyer, D. P.; Lee-Gosselin, A.; Malounda, D.; Swift, M. B.; Maresca, D.; Shapiro, M. G. Acoustic Biosensors for Ultrasound Imaging of Enzyme Activity. Nature Chemical Biology 2020, 16 (9), 988–996. 10.1038/s41589-020-0591-0.

(33) Gyöngy, M.; Arora, M.; Noble, J. A.; Coussios, C. C. Use of Passive Arrays for Characterization and Mapping of Cavitation Activity during Hifu Exposure. In Proceedings - IEEE Ultrasonics Symposium; 2008; pp 871–874. 10.1109/ULTSYM.2008.0210.

(34) Haworth, K. J.; Bader, K. B.; Rich, K. T.; Holland, C. K.; Mast, T. D. Quantitative Frequency-Domain Passive Cavitation Imaging. IEEE Transactions on Ultrasonics, Ferroelectrics, and Frequency Control 2017, 64 (1), 177–191. 10.1109/TUFFC.2016.2620492.

(35) Coviello, C.; Kozick, R.; Choi, J.; Gyöngy, M.; Jensen, C.; Smith, P. P.; Coussios, C.-C. Passive Acoustic Mapping Utilizing Optimal Beamforming in Ultrasound Therapy Monitoring. The Journal of the Acoustical Society of America 2015, 137 (5), 2573. 10.1121/1.4916694.

(36) Gray, M. D.; Coussios, C. C. Broadband Ultrasonic Attenuation Estimation and Compensation With Passive Acoustic Mapping. 2018, 497–516. 10.1109/JPHOT.2016.2538758.

(37) Lakshmanan, A.; Lu, G. J.; Farhadi, A.; Nety, S. P.; Kunth, M.; Lee-Gosselin, A.; Maresca, D.; Bourdeau, R. W.; Yin, M.; Yan, J.; Witte, C.; Malounda, D.; Foster, F. S.; Schröder, L.; Shapiro, M. G. Preparation of Biogenic Gas Vesicle Nanostructures for Use as Contrast Agents for Ultrasound and MRI. Nat Protoc 2017, 12 (10), 2050–2080. 10.1038/nprot.2017.081.

(38) Bader, K. B.; Holland, C. K. Gauging the Likelihood of Stable Cavitation from Ultrasound Contrast Agents. Physics in Medicine and Biology 2013, 58 (1), 127–144. 10.1088/0031-9155/58/1/127.

(39) Collin, J. R. T.; Coussios, C. C. Quantitative Observations of Cavitation Activity in a Viscoelastic Medium. The Journal of the Acoustical Society of America 2011, 130 (5), 3289–3296. 10.1121/1.3626156.

(40) Dutka, P.; Malounda, D.; Metskas, L. A.; Chen, S.; Hurt, R. C.; Lu, G. J.; Jensen, G. J.; Shapiro, M. G. Measuring Gas Vesicle Dimensions by Electron Microscopy. Protein Science 2021, 30 (5), 1081– 1086. 10.1002/pro.4056.

